# Astrocytic engagement of the corticostriatal synaptic cleft is disrupted in a mouse model of Huntington disease

**DOI:** 10.1101/2022.12.06.519168

**Authors:** Carlos Benitez Villanueva, Hans J. T. Stephensen, Rajmund Mokso, Abdellatif Benraiss, Jon Sporring, Steven A. Goldman

## Abstract

Astroglial dysfunction contributes to the pathogenesis of Huntington’s disease (HD), and glial replacement can ameliorate disease course. To establish the topographic relationship of diseased astrocytes to medium spiny neuron (MSN) synapses in HD, we used 2-photon imaging to map the relationship of tRFP-tagged striatal astrocytes and rabies-traced, EGFP-tagged coupled neuronal pairs, in R6/2 HD and wild-type (WT) mice. The tagged, prospectively-identified corticostriatal synapses were then studied by correlated light electron microscopy followed by serial block-face scanning EM, allowing nm scale assessment of synaptic structure in 3D. By this means, we compared the astrocytic engagement of single striatal synapses in HD and WT brains. R6/2 HD astrocytes exhibited constricted domains, with significantly less coverage of mature dendritic spines than WT astrocytes, despite enhanced engagement of immature, thin spines. These data suggest that disease-dependent changes in astroglial engagement and sequestration of MSN synapses enable the high synaptic and extrasynaptic levels of glutamate and K^+^ that underlie the striatal hyperexcitability of HD. As such, these data suggest that astrocytic structural pathology may causally contribute to the synaptic dysfunction and disease phenotype of those neurodegenerative disorders characterized by network overexcitation.

**Significance Statement:** Astrocytic physiological dysfunction contributes to development of the neurodegenerative phenotype in Huntington’s disease (HD), but the structural correlates to this dysfunction are unclear. Here, we used a combination of viral tracing, phenotype-specific tagging, and ultrastructural modalities to reconstruct and study HD synapses at nm scale, in the neostriata of HD mice. We discovered significant impairment in the glial engagement of mature striatal synapses. In light of the known deficiencies in glutamate and potassium uptake by HD astrocytes, these findings suggest the potential for leakage of excitatory synaptic contents during neurotransmission, and hance a structural basis for neuronal hyperexcitability in HD. More broadly, our data suggest that astrocytic structural pathology may causally contribute to those neurodegenerative disorders associated with central hyperexcitability.

## Introduction

Huntington’s disease (HD) is a neurodegenerative disorder in which abnormally long CAG repeat expansions in the first exon of the huntingtin (HTT) gene result in a mutant form of the gene (mHTT), which disrupts glial as well as neuronal physiology, leading to dysfunction first evident in neostriatal medium spiny neurons (MSNs) (1). These cells are the only output neurons of the striatum, and receive the vast majority of cortical and thalamic glutamatergic inputs to the striatum. As such, the MSNs are the integrating units of the striatum, of which they comprise roughly 95% of all neurons. The glutamatergic inputs of these MSNs receive extensive support from astrocytes, via the synaptic clearance of glutamate and potassium ions by astrocytic processes, as well as of modulatory local gliotransmitter release, which together support the homeostatic maintenance of the synaptic environment (2). As a result, the spatial relationship of astrocytic fibers to the individual dendritic spines with which they are associated – and in particular the precise three-dimensional topographical relationship of those fibers to the synaptic cleft - may be critical determinants of synaptic efficiency and firing thresholds (3).

The morphology of a dendritic spine is a determinant of its activation dynamics, and hence of its role and importance within that dendritic network (4). The structural relationship of that spine and its encompassed synapse to the astrocytic fibers with which it is engaged is yet another, critical determinant of its firing threshold, since the astrocytic fibers participate in the sequestration and functional compartmentalization of individual synapses, helping to regulate synaptic activation by removing potassium and glutamate from the synaptic cleft. Yet the structural relationship of synapses with their associated astrocytes has been difficult to study, even moreso to quantitatively describe. Whereas several analytical tools exist for extracting quantitative information from dendritic arbors and spines as captured via fluorescent methods (5–8), analogous tools for structurally rendering nanometer scale astrocytic interactions with individual dendritic spines have not hitherto been available. Instead, recent strategies have focused on the relationship of synapses to their supporting astrocytic elements via both FRET microscopy and super-resolution applications in thick tissue (9, 10). In particular, Khakh and colleagues (9) used FRET in an elegant study of the effect of HD on astrocytic interactions with MSNs, concluding that the involution of striatal astrocytes in HD was associated with some astrocytic retraction from affected corticostriatal synapses. Yet this and analogous studies via FRET provide inferential functional data; they do not provide precise or quantifiable structural characterization of the principal elements of the tripartite synapse – the presynaptic terminal, the postsynaptic dendritic spines, and their enveloping astrocytic fibers. As such, the relationship of mutant HTT-dependent changes in dendritic spine morphology, to disease-associated changes in astrocytic structure, and the effects of these concurrent pathologies on synaptic structure and hence function, has remained unclear. More broadly, such topographic assessment and nanometer-scale reconstruction of perisynaptic environments has remained a challenge, especially so in the adult mammalian brain.

To address this issue, in this study we combined correlated light electron microscopy (CLEM) with serial block-face scanning EM, to directly visualize specific astrocyte-synapse interactions on striatal MSN spines, and then used a novel analytical strategy to extract quantitatively significant information from nanometer-scale structures in these discrete 3D volumes. We combined monosynaptic retrograde tracing using glycoprotein-deleted replication-incompetent rabies virus (11), with astrocyte-specific lentiviral tags to visualize those astrocytes interacting with the targeted synapses. This approach allowed us to define the structural components of synaptic fields in these brains at resolutions of <10 nm, over volumes of 9000 μm^3^ . We chose to study the neuronal phenotype revealed by retrograde tracing of MSNs via their projections to the globus pallidus external segment (GPe), since GPe projection neurons are predominantly D2-expressing MSNs and are the first striatal neurons to manifest dysfunction in HD (12). We found that in HD mice, the synaptic environments of mature dendritic spines experience a marked decrease in both astrocytic engagement and synaptic isolation, suggesting a disease-associated decline in the contribution of astrocytes to both synaptic function and sequestration in the adult brain.

## Results

### Astrocytic engagement of single corticostriatal synapses may be directly visualized

We used correlated light electron microscopy (CLEM) (13) together with 2-channel fluorescence imaging of turboRFP-tagged, retrograde traced MSNs and lentiviral EGFP-tagged astrocytes, to reveal interacting domains of striatal MSN dendrites and astrocytic processes in the adult mouse neostriatum (**Figure 1**). This combination of imaging modalities allows the capture of relevant light microscopy data prior to serial EM scans, facilitating retrieval of the desired regions of interest (ROI) in EM, and preserving contextual information of the nanometer-scale, serial SEM datasets (**Figure 1C, supplementary figure 1**). Using this approach, we first captured EM datasets of specific MSN dendritic spines displaying astrocytic processes in close proximity to the periphery of the post-synaptic density (PSD). We followed this with a novel quantification strategy to assess the global spatial engagement of astrocytic processes cradling the synaptic environment, and the effects of HD on these structures.

**Figure 1.**
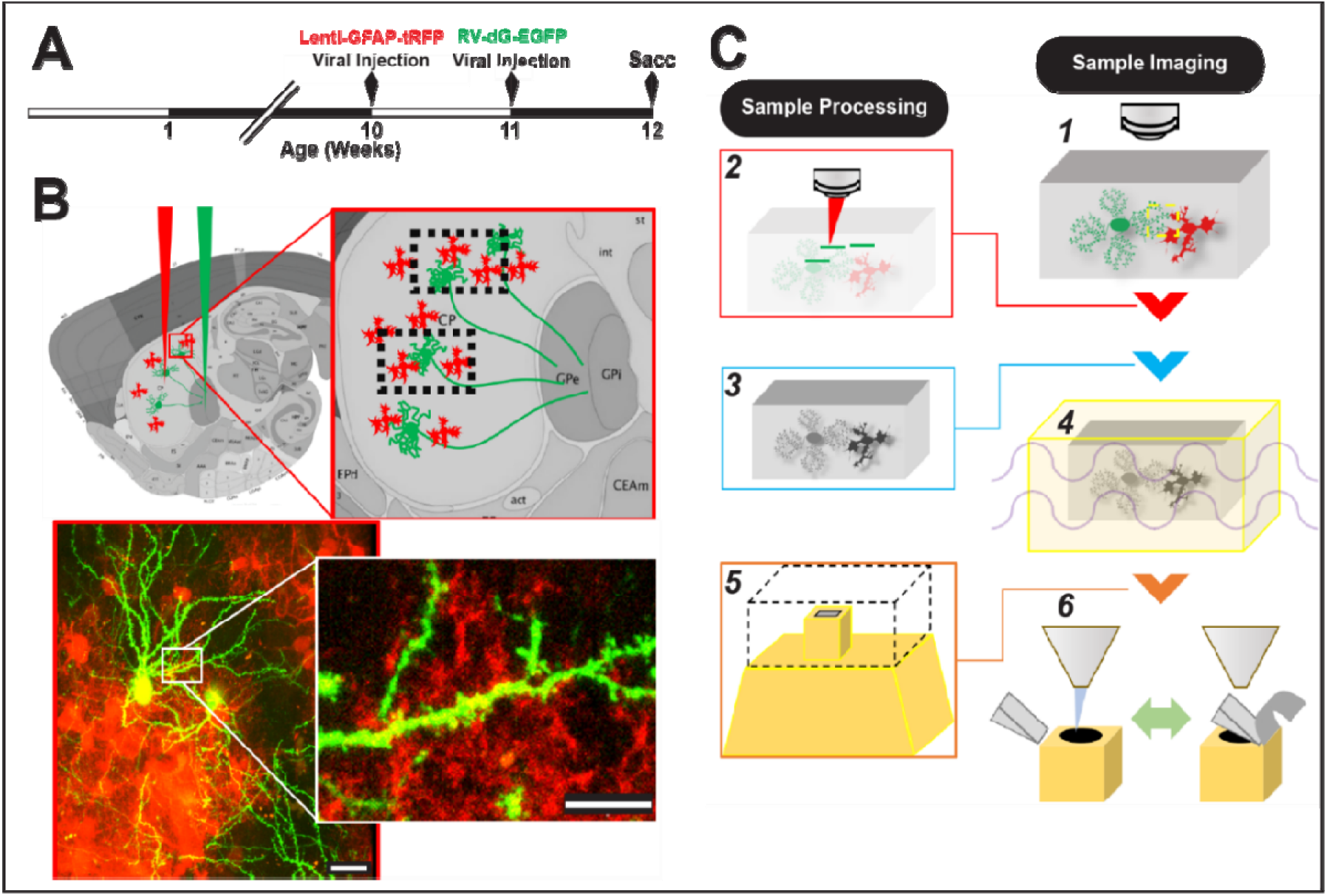
Phenotype-specific ultrastructural analysis of synaptic-glial interactions using SBF SEM. Rabies tagging of neuronal pairs was combined with lentiviral GFAP-GFP-based identification of coupled astrocytes, as assessed by correlated light EM (CLEM) and serial blockface scanning electron microscopy (SBF SEM), to describe prospectively-defined astrocyte-engaged synaptic fields. **A**. Timeline of sequential viral deliveries. **B**. Astrocyte-engaged synaptic fields identified by pallidal Rabies retrograde tagging of MSN neuronal network combined with astrocytes-specific Lenti-GFAP-tRFP. **C**. Workflow by which sequential multimodal imaging is used to pinpoint ROI: 1) 2-photon imaging is used to identify interacting domains; 2) 2-photon branding is used to induce nearinfrared ablations in the vicinity of the ROI; 3) EM sample preparation via dehydration, osmium impregnation and resin embedding; 4) MicroCT scan of entire sample; 5) Relocalization of ROI via microCT cross-reference, and trimming of excess material; 6) Serial block-face SEM of ROI domain. Scale: 10 μm

To that end, we injected GFAP-tRFP lentivirus bilaterally into the striata of R6/2 x rag1 ^-/-^or WT rag1^-/-^ control mice at 10 weeks of age, followed a week later by intrapallidal injection of replication-inefficient rabies virus with EGFP substituted g-protein (RV-dG-EGFP) (**Figure 1A**). A week thereafter, at 12 weeks of age, we killed the mice and used correlated light EM (CLEM) to isolate synaptic ROIs, focusing on regions with pre-defined astrocytic processes intermingling with dendritic arbors of MSNs. These regions were identified, ablated, and selectively isolated through correlative methods (**Figure 1B**), culminating in serial SEM acquisition for nanometer-scale data acquisition (**Figure 1C, Figure S1; Video 1**). Data were collected from striatal sections sampled from 2 WT and 3 R6/2 animals. We focused on dendritic branches of at least 3^rd^ order, and only dendritic spines that displayed astrocytic engagement; only those spines engaged by astrocytes were included in this analysis. Of the 144 total astrocyte-synapse ROIs we selected, 22 did not present astrocytic apposition within the first 100nm of the radius surrounding the PSD once the SEM series were segmented, and thus were not included in this analysis. The remaining 81% of the ROIs (117 astrocyte-synapse ROIs) were analyzed for various metrics, indicating a modest success rate for capturing specific astrocyte-synapse interactions exhibiting neuron-glia engagement.

We observed a variety of structural configurations with regards to relationships of astrocytic processes engaging synapses, both pre- and post-synaptically. To standardize our quantitative descriptions of these synapses, we focused on their post-synaptic densities. We identified the post-synaptic densities as regions on the head of the dendritic spine presenting high electron dense material in the EM datasets, as expected for asymmetric glutamatergic synapses. Once we identified prospective ROIs, defined as regions within which astrocytic domains revealed by tRFP engaged EGFP-expressing dendritic spines of striatal MSNs, we imaged those volumes containing the selected ROIs. We then selectively ablated points within that ROI with high power IR laser scans, so as to generate auto-fluorescent, spatially discrete fiducial features (**Video 2**) (14). From these ROIs, we traced specific segments across 3 orthogonal imaging scales (2-photon, x-ray tomography, and serial EM), and then correlated the corresponding datasets so as to acquire the tripartite synapses found within the original ROI (**Video 2**). By this means, we were able to directly visualize our prospectively-defined synapses of interest.

### Astrocytic proximity and peripheral coverage of synapses is disrupted in R6/2

We next postulated that for regions of interacting astrocyte processes and MSN dendritic domains, that the degree of infiltration of the synaptic periphery by the astrocyte would be affected by HD-associated pathology. To test this postulate, we re-localized our regions of interest, identified the corresponding dendritic domain, and then manually segmented those spines engaged by astrocytic processes (**Figures 2A-B**). We generated regions of analysis that incorporated features derived from the PSD, modeling that region on a torus structure surrounding the synaptic cleft (**Figure 2C**). We set the bounds of analysis as a torus-shaped region surrounding each PSD, so as to normalize for the 3-dimensional variability of these structures within and across datasets. Specifically, we measured the longest axis of the PSD structure, and defined the inner and outer bounds of the virtual torus surrounding the PSD as 0.75 * R_max_ and 1.5 * R_max_, respectively, so as to establish a consistent link between the region analyzed and PSD morphology. The depth of the torus was set to 300 nm, as the thickness of each PSD was consistent, regardless of the PSD area on the dendritic spine head; the center of the virtual torus was set to the 3D center the PSD. Anything outside of this torus was excluded and we only captured astrocytic data only within the torus-like structure for quantification. By this approach, we defined the peripheral limits of analysis, while normalizing the variability across PSD configurations (**Figures 2D-E**); we quantified the volume of the PSD itself, the minimum distance between the PSD edge and the astrocyte membrane, and the total volume of astrocytic cytoplasm for each synapse within each segmented synaptic environment (**Figure 2 F-H**).

**Figure 2.**
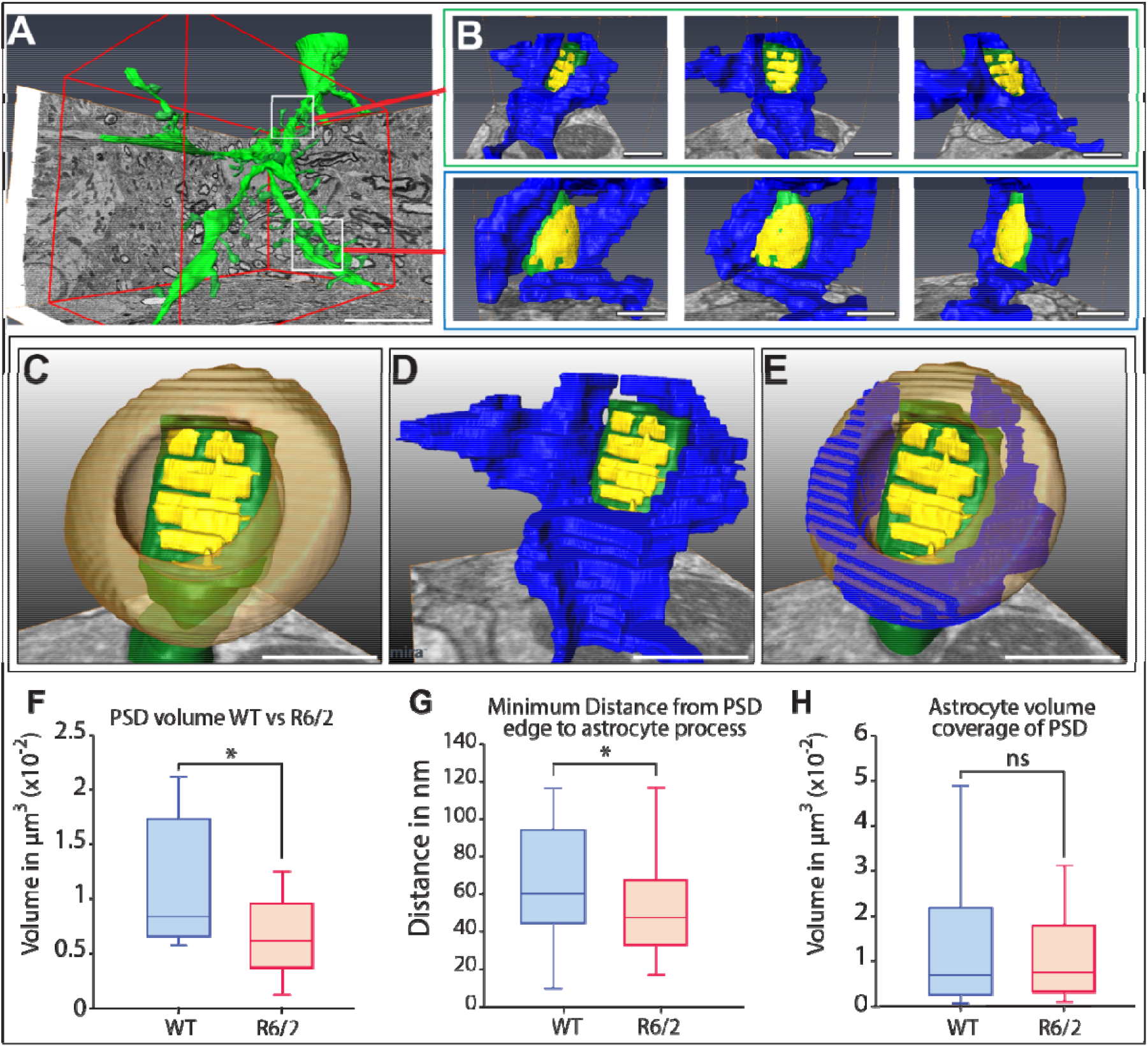
Analysis of astrocyte-synapse interactions. **A.** Detailed view of dendritic branch and spine reconstruction**. B.** Reconstruction of selected spines and perisynaptic astrocytic domains in 3D, with 3 views around axis for each spine (spines, *green;* post-synaptic densities, *yellow;* peripheral astrocyte domains, *blue*). **C.** Definition of analytic boundaries of torus region parallel to the PSD, with outer bounds defined as: R_outer_bound = R_max_ (of PSD) * 1.5 and R_inner_bound = R_max_ * 0.75. Depth/thickness of torus, 300 nm. **D.** Astrocyte domain contained within the periphery of the PSD (blue, transparent). **E.** Exclusion of excess material contained outside the bounds of analytical torus and highlight of astrocyte domain (*blue*, opaque) within the total analysis domain (*orange*, transparent). **F-H.** All data were limited to the pre-defined analytic domains. **F.** Volume of post-synaptic density (nonequal variances, t-test two tail, *p=0.03). **G.** Minimum distance from PSD edge to astrocyte process edge, non-equal variances (F-test 0.009, t-test *p=0.04). **H.** Astrocyte coverage within a radius 1.5 * r_max_ of PSD. Equal variances, non-significant (**p = 0.324, n.s.**) (WT: 2 mice, n=32; R6/2: 3 mice, n=45). Scale: 5μm.

We found the volume of the segmented PSD exhibited a significant decrease in R6/2 mice relative to WT, with an average of 10.9 ± 1.6 x 10^-3^ μm^3^ for WT and 6.3 ± 0.9 x 10^-3^ μm^3^ for R6/2, spanning a range of 5.6 - 21.1 x 10^-3^ μm^3^ for WT and 1.2 - 12.8 x 10^-3^ μm^3^ for R6/2 (**Figure 2F**; p <0.05, two tailed t-test; WT, n = 32 synapses, R6/2, n = 43). We also found the minimum distance from the PSD boundary to the astrocytic process edge to be significantly reduced in R6/2 synapses - indicating closer proximity - with an average distance of 73.5 ± 8.1 nm for WT vs. 58.7 ± 4.5 nm for R6/2 (p<0.05, WT: n=32 synapses; R6/2: n=45 synapses**; Figure 2G**). However, despite an apparent reduction in the astrocytic coverage of the synaptic periphery in R6/2 mice, this initial approach did not reveal an overall reduction in peri-synaptic astrocytic volume in R6/2 mice, relative to their WT controls (**Figure 2H,** p>0.05). Rather, this initial analysis indicated that HD R6/2 mice exhibit diminished PSD size, with closer apposition of astrocytic processes to the PSD. In that, these data support the aforementioned FRET study of Khakh and colleagues (9), which observed closer apposition of astrocytic processes to striatal synapses in HD mice than in WT controls. To resolve this apparent paradox of closer astrocytic proximity yet diminished areal coverage of R6/2 synapses, we next sought to parse out the detailed topographies of HD and WT MSN synapses, focusing on both the detailed astrocytic geometry at their respective synaptic clefts, and on the types of synapses - and hence the structural maturity of the dendritic spines - with which those astrocytes are engaged.

### Astrocytic infiltration of the synapse is altered as a function of both HD and spine class

To thus describe the structural relationships between individual synapses and their partnered astrocytes, we established a model by which the topographies of individual synapses and their astrocytes could be accurately rendered and quantitatively described. For each modeled synapse, we measured the astrocytes in a progressively larger region around the PSD. These extended regions can be understood as a dilation or inflations of the PSD, or more formally, as the set of all points within a progressively increasing distance from the PSD edge. This forms a set of measurement functions defining geometrical features of the astrocyte as a function of the distance to the PSD. As such, each synapse contributes a set of measurement functions which, taken together, can be understood as a cross-K-function summary statistic (15). By this means, we analyzed the interactions of the astrocytic processes as they intersected the virtual expansions of the PSD. In particular, this measurement method allowed us to estimate the astrocyte volume and surface area as a function of the distance to the PSD (**Figure 3**) (16). By this approach, as the PSD is dilated, the points and characteristics of the dilation’s interaction with the surrounding astrocyte are recorded.

**Figure 3.**
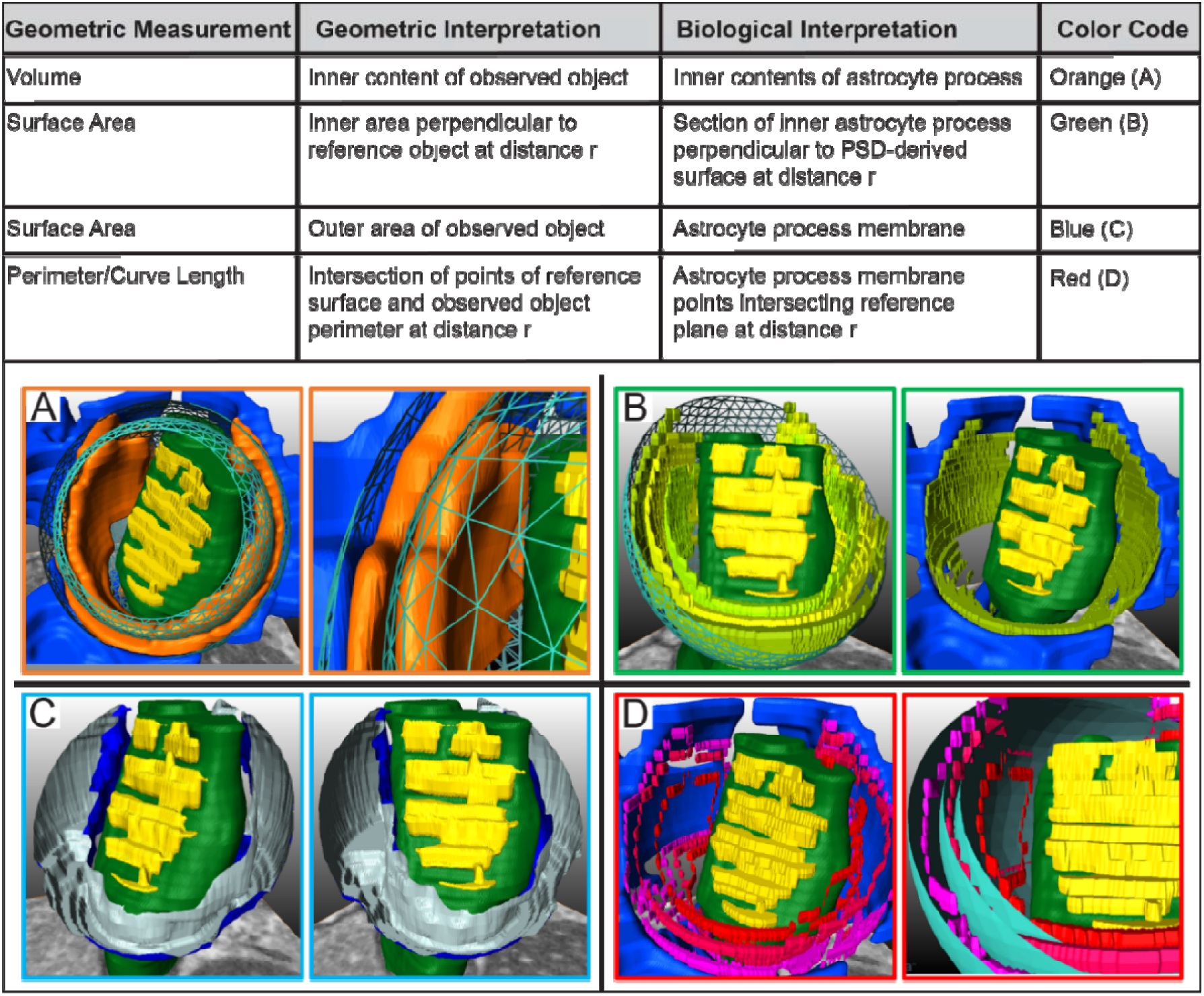
Modeling the topographic relationship of astrocytes to dendrites at the synapse. To model the topology of astrocytic synaptic engagement, we used a spherical approximation of geometric measurements, defined as at the intersections of interacting reference and observed surfaces, at increasing distance (radius) from the PSD. To this end, we assessed the 3D interactions of the astrocytic processes with the synapse, and derived geometric measurements (Hausdorff measures) for each astrocyte process (observed object). These geometric derivations included: Astrocyte surface area (in *blue*), volume (*orange*), surface area perpendicular to reference object (*green*), and intersection points (contour), in *red*. **A.** Representative view of astrocyte volume surrounding the PSD (*orange*, astrocytic cytoplasm) measured over discrete measurements represented by teal intervals surrounding the PSD. **A,** *left panel*, overview; **A,** *right panel*, identification of discrete intervals and quantified material. **B.** Discrete measurements of astrocyte surface area perpendicular to the PSD (*left*) shown at increasing radii intervals from PSD. Excluded astrocyte material shown in blue (*right*) **C.** Surface area of astrocyte surrounding PSD within the pre-defined analysis limits (*blue*, corresponding to cell membrane). View from 2 angles. **D.** Contours where astrocyte structure intersects PSD-derived surface. Multiple iterations of contours at intervals (**D**, *left* panel). Contour overlay shows measurement steps in *teal* (**D**, *right*).

Past studies of astrocytic calcium dynamics have reported the derivative of the astrocytic volume to be highest within 200 nm of the post-synaptic densities (17). In addition, simulations of extra-synaptic glutamate receptor (GluN) diffusion by Gavrilov and colleagues have suggested that this distance covers the glutamatergic activity profiles important to LTP and LDP dynamics (18). On that basis, we chose to focus on, and analyzed, the geometric characteristics of the astrocyte processes within a 150 nm range spanning 25-175 nm of the PSD.

A number of studies have reported that astrocytic coverage of central synapses is linked to spine maturity (19–22). We therefore categorized our observations on astrocytic interactions with synapses on the basis of dendritic spine morphology and inferred maturity (**Video 3**). In particular, we assessed the astrocytic engagement of separately identified subpopulations of thin and mushroom spines, which roughly correspond to immature and more mature dendritic spines (23–26). We separately validated these different categories of spines utilizing k cross-section statistical methodology (**Figure S3**) (16). Using this analytic strategy, we then asked if the apposed astrocytic volume associated with each synapse was significantly altered in HD mice, and if this varied as a function of spine maturity.

To this end, we first measured the astrocyte volume in the periphery of the synapse, and found significant atrophy on mature, mushroom-like synapses (**Figures 4A**). In particular, mushroom spine synapses displayed a significant decrease of engaged astrocytic volume, both in cumulative terms and upon distance-normalized analysis (**Figures 4A** and **S4A,** *bottom panel*). In contrast, we found a trend towards increased astrocytic volume on thin spine synapses, both in their cumulative or distance-normalized volumetrics (**Figures 4A** and **S4A,** *middle panel*), indicating the R6/2 HD striata experiences an over-sequestration of the peri-synaptic environment, relative to WT. Again analyzing within 175 nm of the PSD edge, we found that the average volume of astrocyte coverage of mushroom spines in R6/2 was 7.4 ± 0.17 x 10^-4^ μm^3^, vs 13.7 ± 0.2 10^-4^ μm^3^ for WT. In contrast, R6/2 thin spines displayed astrocyte volumetric engagement of 10.6 ± 3.3 x 10^-4^ μm^3^, vs 6.9 ± 1.3 10^-4^ μm^3^ for WT controls. Together, these data indicate that mutant HTT expression in R6/2 mice is associated with a substantial diminution of astrocytic apposition to synapses on mature, spines of striatal MSNs, but no such loss of astrocytic engagement with immature, thin spines within the same striata.

**Figure 4.**
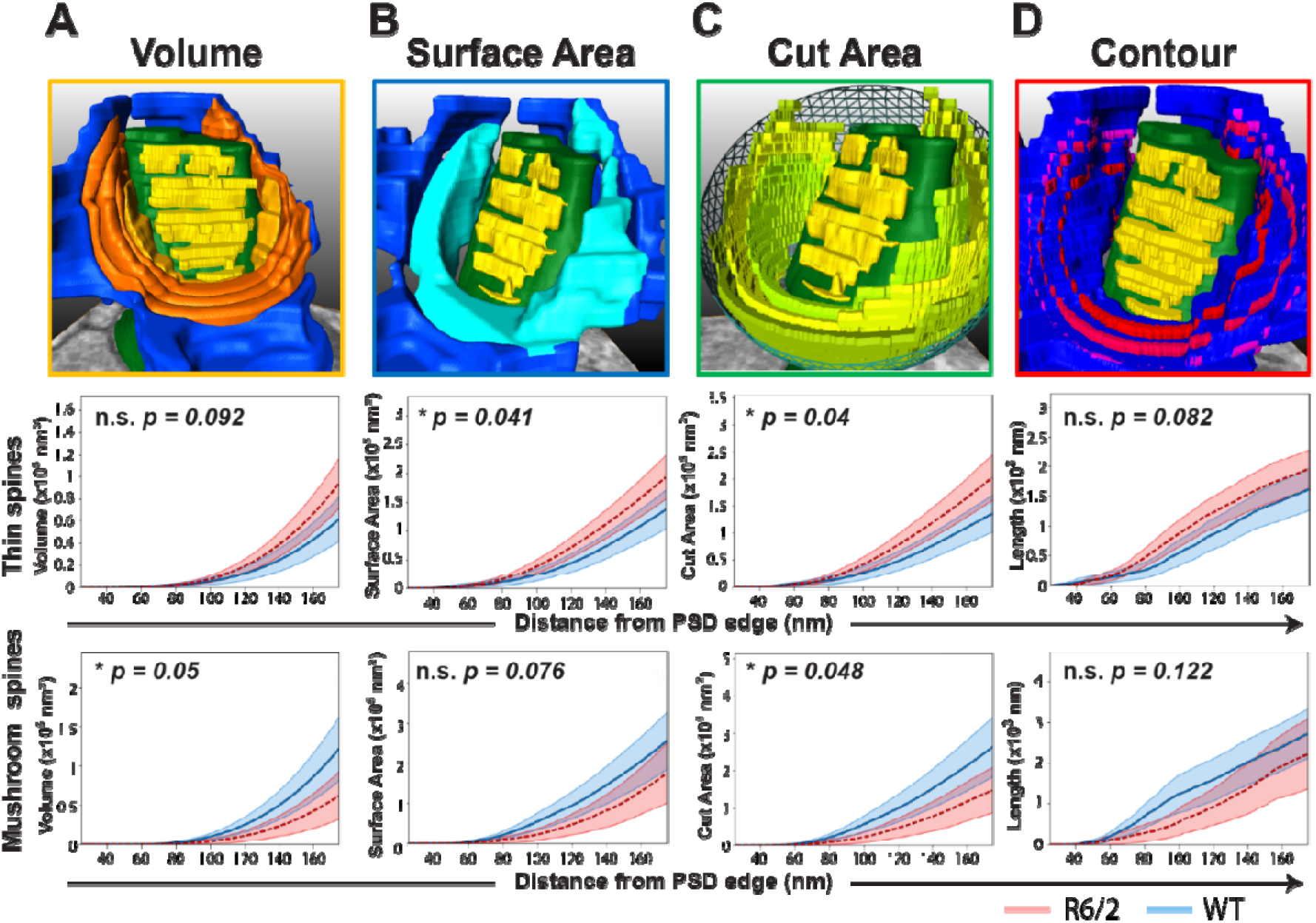
HD affects the geometry of astrocyte-synapse interactions. Analysis of astrocyte-synapse interactions using a derivation of k cross-section statistics. Perisynaptic astrocytic volume, surface area, perpendicular area, and contour length, were each measured over the proximal interval of 25-175nm from the PSD edge; cumulative data shown. **A. Perisynaptic astrocytic volume** Thin/immature spines, with trend to increased associated astrocyte volume in R6/2 relative to WT (*top panel*, p=0.09, n.s.). Mature/mushroom spines, with significantly decreased associated astrocytic volume (*bottom*, *p=0.05). **B. Surface area** Thin immature spines of R6/2 mice were engaged by an increased surface area of their associated astrocytes (*top panel*, *p=0.04). In contrast, the neighboring astrocytes of mature/mushroom spines, trended to present a decreased surface area to the synapse in R6/2 mice (*bottom*, p=0.08, n.s.). **C. Cut area** Thin/immature spines, increased neighboring astrocyte perpendicular surface area (*top panel*, *p=0.04). Mature/mushroom spines, decreased neighboring astrocyte orthogonal surface area (*bottom*, *p=0.048). **D. Contour** Thin/immature spines, unchanged neighboring astrocyte contour length (*middle panel*, p=0.082, n.s.). Mature/mushroom spines, unchanged neighboring astrocyte contour length: (bottom panel, p=0.12, n.s.) Thin/Immature spines: WT, 4 mice, n=30 spines; R6/2, 5 mice, n=35 spines. Mature/mushroom spines: WT, 4 mice, n=27 spines; R6/2, 5 mice, n=22 spines. Monte Carlo permutation method, absolute AUC, 95% CI shown

### Peri-synaptic astrocytic surface area is altered in HD differentially by dendritic spine type

We next asked if R6/2 astrocyte processes exhibited disease-dependent changes in their peri-synaptic surface area, and whether this too might vary as a function of spine class. To this end, we analyzed our structural data as a function of both the distance from the PSD boundary, and of spine class (**Video 4**). We found that mushroom spine synapses exhibited a strong but less than significant trend towards a diminished astrocytic surface area cradling the synaptic cleft, as might be predicted on the basis of the sharply decreased astrocytic volume associated with these synapses (p= 0.076) (**Figures 4B** and **S4B,** *bottom panel*). In contrast though, we found that the surface area of astrocytic processes engaging thin spines was increased in R6/2 MSNs, relative to controls (p = 0.041) (**Figures 4B** and **S4B,** *middle panel*). In particular, the average surface area of astrocyte coverage upon mushroom spines (within 175 nm of the PSD edge) in R6/2 was 1.93 ± 0.47 x 10^-5^ μm^2^, vs 2.75 ± 0.53 x 10^-5^ μm^2^ for WT. In 5 contrast, R6/2 thin spines encountered peri-synaptic astrocyte surface areas of 2.08 ± 0.66 x 10 μm vs 1.5 ± 0.27 x 10 μm for WT mice. Overall, these results indicate differential effects of mHTT expression on astrocytic membrane geometry and engagement on thin and mushroom spines, with the loss in R6/2 mice of astrocytic membrane availability within the vicinity of mature, mushroom spine synapses, despite the concurrent increased astroglial coverage of thin spines, as assessed by surface area.

Among the derivations from the cross-K statistic methodology, we also quantified the cut area and contour extension of the intersecting objects (i.e. dilated PSD reference object and astrocytic process). While surface area and volume are metrics with clear biological identities, perpendicular area and contour are derived functions. Nevertheless, we found the astrocyte perpendicularly cut area to be significantly decreased for mushroom spines (**Figure 4C,** *bottom panel*), and increased for thin spines (**Figure 4C,** *middle panel*). Conversely, the contour measurements displayed no significant difference when we compared the astrocytic coverage of thin vs mushroom spines (**Figures 4D**).

Over the perisynaptic distance interval studied, we were presented with the difficulty that cumulative comparisons and differences can be disproportionately inflated as a function of increasing radius. Much like the area of a circle increases proportional to the square of the radius, the surface area of a spherical domain of interest increases by the cube of the radius from the PSD. This exponential increase in surface area can be controlled for by normalizing the data, which considers the recorded cumulative information in comparison with the exponential increases per radius step. This normalization is analogous to converting the total astrocyte coverage to proportional coverage. When these data were normalized to radial distance from the PSD, the disease (R6/2 vs WT)-dependent atrophy of astrocytic processes associated with mushroom spines was sustained across all geometric metrics (**Figure S4**).

Of note, our classification of thin and mushroom spines was verified using the same method of k cross-section statistics, setting the observed object as the spine rather than its associated astrocyte. By this means, we found that while the surface area and volume of the two spine classes – thin and mushroom – differed from one another, they did not differ significantly between WT and R6/2 mice (**Figure S3**).

## Discussion

In this study, we developed a methodology for quantitatively assessing the structural relationships of astrocytes to their partnered synapses, and for doing so as a function of glial disease, we designed two complementary approaches to generate spatial information sufficient to describe the geometry of synaptic environments. The first involved the modeling of a torus region, defined by the PSD size, over which several relationships between the PSD and the astrocyte could be quantified. The second strategy used k cross-section statistics to derive the 3D geometric qualities of the astrocytic engagement into discrete, quantifiable parameters. This new imagebased quantitative analytic protocol provides us a new means of assessing and quantifying the structural correlates to both normal and pathological synaptic function. In this study, we employed this methodology to assess the effects of Huntington’s disease on astrocytic engagement of striatal synapse.

Using this strategy, we found that astrocytic engagement of corticostriatal excitatory synapses on medium spiny neurons is significantly impaired in the R6/2 mouse model of HD. This disease-associated defect in glial synaptic engagement was manifested by abnormalities in astrocytic sequestration of the post-synaptic density, as reflected by astrocytic retraction from mature dendritic spines, despite paradoxically increased astrocytic apposition to thin, immature spines. These data suggest that while the astrocytic engagement of glutamatergic synapses varies with spine maturity, the structural relationship of astrocytes to their partnered synapses in the HD striatum is abnormal at multiple stages of spine maturation.

Several recent studies of HD mice have noted a disease-dependent reduction in astrocytic domain size and in the number of close proximity synapse-astrocyte sites (9, 27). In the present study, we sought to more precisely define the effects of HD on synaptic ultrastructure, by combining rabies viral tagging of defined synaptic pairs, and lentiviral-EGFP tagging of their partnered astrocytes, with electron microscopy to establish a correlative light-electron microscopy (CLEM)-based approach to study the astrocyte-synapse interactions between specific, prospectively-identified pre- and postsynaptic neuronal partners. In particular, we combined rabies tagging of corticospinal-MSN synapses with lentiviral-GFAP-driven EGFP identification of local astrocytes, and examined the resultant tissue by sequential imaging using 2-photon, X-ray, CLEM and serial bloc-face SEM (SBF). By this means, we were able to investigate the structural topography of corticospinal-MSN synapses together with their partnered astrocytes with a never-before achieved level of resolution and specificity. In thereby resolving the synaptic environment, we were surprised to observe that these different spine morphologies were associated with such different degrees of astrocytic coverage. This was unexpected, as recent studies had suggested that spine size per se does not determine the astrocytic volume fraction in the synaptic microenvironment (18). Nonetheless, we found that HD astrocytes exhibited increased apposition to thin, nominally immature spines, while more mature, mushroom-like spines enjoyed significantly less astrocytic engagement. As such, these data indicated that the relationship of astrocytes to synapses of different spine maturational classes varies with the disease environment (**Figure 5**).

**Figure 5.**
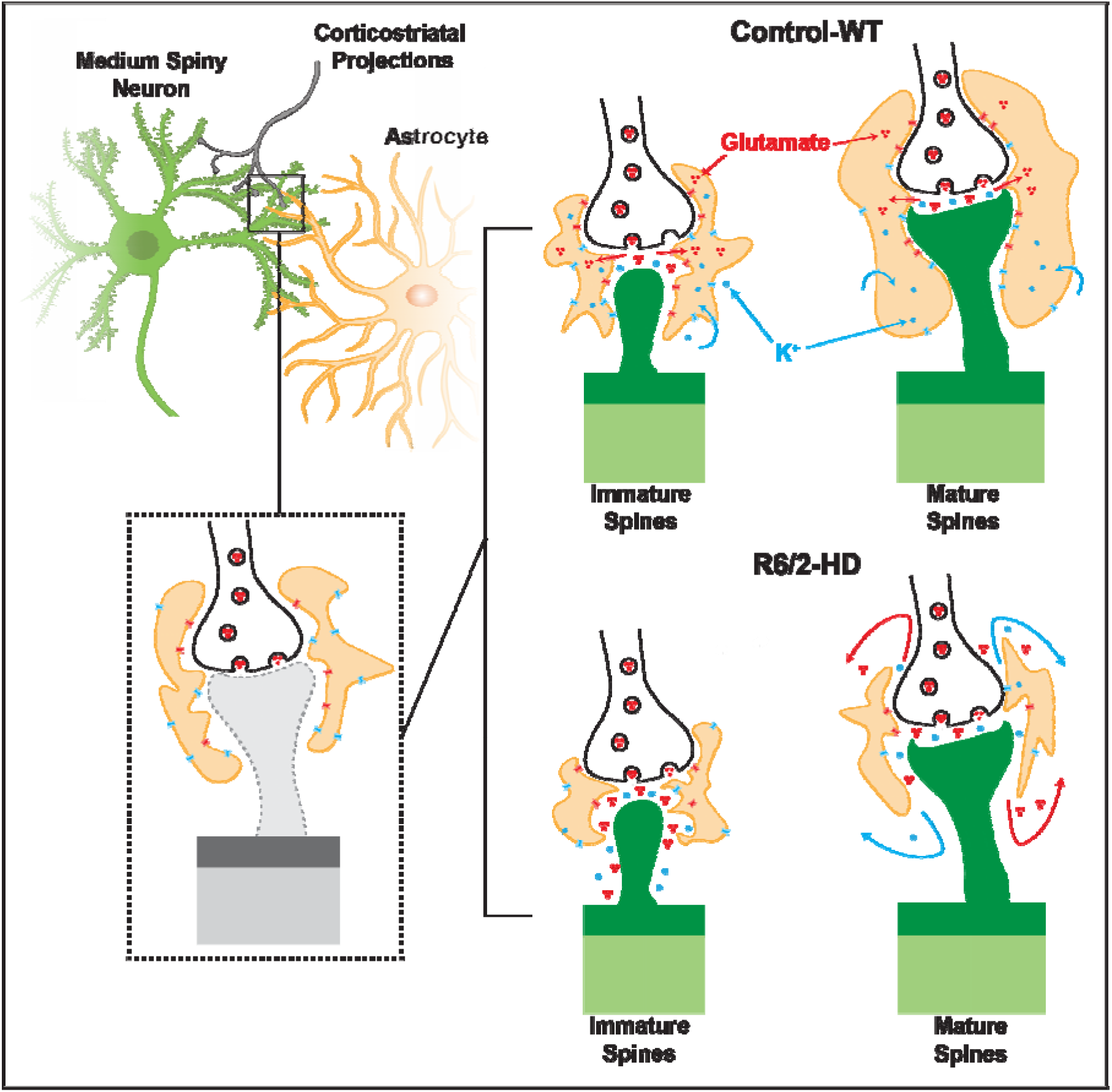
Changes in the peripheral astrocytic coverage of tri-partite dendritic spines of R6/2 mice. Thin/immature spines of young adult mice display increased engagement of synaptic periphery in R6/2 mice, increasing sequestration of glutamate and ions. Mature spines however, experience significant retraction of astrocytic processes from the synaptic periphery, exacerbating the leakage of K^+^ and Ca2^+^ ions, as well as glutamate, which contribute to nonspecific signaling and activation of extra-synaptic receptors.

In this regard, we found that while mHTT-expressing astrocytes maintained their engagement with immature spines, the effects of astrocytic involution were more profound on those resident neurons with more mature spines, which suffered both astrocytic retraction from, and de-sequestration of, their embedded synapses. Indeed, the maturity-dependent decline in astrocytic engagement of striatal synaptic spines, despite retained glial fiber proximity, suggests that R6/2 astrocytes become atrophic, and that this process becomes most manifest in the most mature synapses, identified by the mushroom-like morphologies of their postsynaptic spines. As such, their distal processes might maintain proximity, but more broadly, they lose volume within the peri-synaptic region; as a result, the mature spine synapse loses its astrocytic investment, and hence its spatial sequestration. In contrast, the close relationship of astrocytes to synapses along thin R6/2 spines appears to reflect a sustained, if not actually tighter, engagement of the synaptic periphery by R6/2 astrocytic processes. Yet since HD glia are deficient in K^+^ and glutamate uptake (Osipovitch et al., 2019), these thin spine synapses might be expected to suffer increased intra-synaptic K^+^ and glutamate levels even with sustained spatial sequestration (**Figure 5**), hence contributing further to the hyperexcitability of striatal neuronal networks in HD. Overall, the abnormal patterns of astrocytic engagement with both mature and immature spines, and their attendant alterations in the peri-synaptic sequestration of both neurotransmitters and neuroactive ions, would be expected to degrade input-specific responses, confounding LTP and LDP dynamics at the cellular level (17, 18), while disrupting network coordination at the systems level.

Although the highly branched, bushy structure of astroglial processes makes the astrocyte a difficult domain to study structurally, its functional role in the maintenance of synaptic homeostasis can be leveraged to infer fine morphological changes. In that regard, a recent study by Khakh and colleagues exploited the phenomenon of Förster resonance energy transfer (FRET) to study the points of closest proximity of astrocytic processes to synapses (9). To do so, they selected and modified presynaptic- and peri-synaptic components to express mCherry and GFP, respectively. They then induced fluorescence by one fluorophore and recorded emission events from the other, capitalizing upon the occurrence of two-color emissions only in those regions where the fluorophores are in close enough proximity for FRET events. Yet while elegant, FRET events, at 50% efficiency, only occur at those sites where GFP and mCherry localize within 5.1 nm of each other; in general, FRET is sensitive to changes over 10 nm, where efficiency ranges from 5% to 95%. Thus, while this method reveals the existence of close-proximity interactions between astrocytic and neuronal structures, it is a semi-binary readout, providing information primarily on the existence – or absence - of synapse-astrocyte interactions within 10 nm of each other. FRET does not provide information on the characteristics of these interactions on an individual spine-by-spine approach, nor does it provide information regarding the astrocyte interaction beyond 10 nm. In similar studies by Gavrilov et al., analyzing astrocytic coverage of synapses in the hippocampus expanded the analysis volume to 600nm from the PSD. According to their post-synaptic receptor activation simulations, the open probability of extra-synaptic GluA receptors peaks at r = 0.15μm (from PSD edge), while GluN receptors seemed to display a generally higher peak than GluA but decayed more steeply than GluA (18). These data suggested that the region of astrocytic engagement most relevant to synaptic function lay within ~200 nm of the PSD; as such, we chose to investigate the geometries of peri-synaptic astrocytic processes within that range.

Recent studies have highlighted the contribution of glial pathology to Huntington’s disease (9, 27–34). These studies have pointed out that mHTT-associated astrocytic dysfunction in particular may have widespread consequences on neuronal as well as glial function. These effects include a disruption in synaptic K^+^ uptake (27), with insufficient clearance at synapses (35); neuronal hyperexcitability due to high intrasynaptic glutamate as well as K^+^, with attendant excitotoxicity (36); and an overall reduction of the astrocytic domain size, and hence in synaptic coverage (9, 27, 28, 35, 37). Parallel studies have highlighted the relationship of astrocytic engagement to spine size and maturity (18, 22, 38, 39); this issue is of particular relevance here due to the scaling of AMPA receptor content to PSD size, and therefore to spine maturity (40–44). In the present study, we extended these observations by comparing the astrocytic engagement of striatal synapses in HD transgenic and wild-type brains, focusing on the effect of HD on the relationships of perisynaptic astrocytes and their processes to single striatal synapses.

As in our structure-based study of synaptic topologies in HD, Khakh and colleagues’ recent study using FRET to assess the proximity of astrocytic processes to synapses found an overall reduction in the R6/2 HD striata in the volumes of astrocytic domains, despite the closer proximity of those glial-synaptic interactions that did occur (9). Yet such measures tell an incomplete story: R6/2 striatal MSNs experience a significant reduction in both their dendritic arbors and in the density of their dendritic spines (12). Furthermore, the proportion of immature to mature spines on the remaining dendrites of 12-week-old R6/2 MSNs is markedly increased, suggesting that most of the functional spines remaining in these mice are thin and relatively immature (45). Together, these data suggest that FRET readouts from the R6/2 HD striatum may derive predominantly from thin spines, which – unlike mature spines – may manifest closer astrocytic engagement, which appears transient and may be compensatory, but which is any case lost with spine maturation. As such, the reduced glial engagement of mature R6/2 synapses, while still clear and significant in our data, may nonetheless be artifactually minimized in our overall data set, owing to the relative under-representation of mature synapses in the late-stage R6/2 striatum (45, 46).

There is mounting evidence that morphological changes in neuroglial interactions are both malleable and activity dependent. For example, the induction of LTP on hippocampal spine terminals leads to the mobilization of neighboring astrocyte processes within 5 min of LTP induction (47). Other recent studies have highlighted the intricate set of signals between neurons and glia that cooperatively regulate synapse formation and maturation (3). While morphological changes per se comprise only one aspect of the changes occurring within these complex environments, these geometric readouts reveal critical information regarding the functions and stability of synapses in response to disease. Yet querying the structural interactions between neuronal spines and glial processes has proven technical challenging. Mouse protoplasmic astrocytes contain tens of thousands of spines within their individual domains, and human astrocytes may contain 1-2 orders of magnitude more synapses than their murine counterparts (48). The diffuse web-like nature of astrocytic processes within the neuropil complicates their study further, as the cells extend processes into the smallest reaches of the extracellular space. These astrocytic domains and their encompassed synapses are difficult to probe in detail using light-microscopy, and yet at the same time are difficult to associate with phenotype using serial EM. Overcoming these obstacles, we found that glial-synaptic interactions within these domains can indeed be defined, captured, imaged at nm scale and analyzed topologically, using the serial combination of lentiviral astrocytic tagging, rabies tagging of coupled neuronal pairs, multiphoton imaging with laser and X-ray registration, followed by serial blockface EM and correlated light EM, that we have outlined.

Using this approach, we found significant structural abnormalities in the astrocytic engagement and investiture of striatal synapses in R6/2 mouse model of HD. In particular, we noted a pattern of diminished engagement of mature synapses by perisynaptic astrocytes, which suggested the diminished sequestration of those synapses, by glia already deficient in K^+^ and glutamate uptake mechanisms. Such disease-associated synaptic pathology may result in the local spread of synaptic glutamate and K^+^ into adjacent synaptic fields, and thereby contribute to the high interstitial K^+^ and attendant hyperexcitability of the R6/2 striatum. More broadly though, the methodologic pipeline that we have outlined in this study allows for the topological investigation of glial-synaptic relationships within any parenchymal domain of interest whose pre- and postsynaptic components are known, and for which associated glial components may be prospectively tagged – whether they are of astrocytic, microglial or progenitor phenotype, and whether of host or exogenous origin. As such, we expect this approach to be broadly useful in defining synaptic microstructure in the neurodegenerative and neuropsychiatric disorders, providing nanometer-scale insight into the integrity of involved synapses in disease states, and going forward, of the structural plasticity of these synapses in response to therapeutic manipulation.

## Materials and Methods

### Animals

All experiments were approved by the Animal Ethics Committee of the University of Copenhagen (Afdeling for Ekperimentel Medicin, AEM Project Plan P-20-054), as well as by the Institutional Animal Care and Use Committee of the University of Rochester. Wild-type females with ovary transplants from R6/2^+^ (120 +/- 5 CAG) donor mice were purchased from Jackson Laboratories (Bar Harbor, ME). The mice were genotyped following weaning and double heterozygous mice were further analyzed to determine their CAG repeat number through PCR with primers encoding a product spanning the repeat region as previously described (49). All experiments included 4-5 mice/group, as indicated, with equal numbers of males and females.

### Induction of fluorescence

Rag1-/- and R6/2 mice were given two viral injections to induce EGFP and tRFP expression in MSNs and Astrocytes respectively. The mice were anaesthetized with isofluorane flow and treated with Xylocain was used as local anaesthetic (0.2mg/mL, delivered as needed but kept below 0.7mg/kg) by inserting the needle under the scalp and retracting the tip as the solution was delivered. At 10 wks of age, the mice received 500 nL bilateral injections of Lenti-GFAP-tRFP in the striatum at the following coordinates: AP +0.5 mm, ML ± 2.0 mm, DV −3.4 mm from Bregma. After surgery, the mice were observed during waking and given carbprofen for pain management (NSAID, 1 mg/mL or 0.1 mL for a 20g mouse). At 11 wks of age, the same operative procedures were followed, but this time with bilateral delivery of replication inefficient retrograde rabies virus RV-dG-EGFP in the Globus Pallidus (coordinates: AP −0.3 mm, ML ± 1.7 mm, DV −3.5 mm) (**Figure 1A-B**) (11).

One week later the animals were transcardially perfused with 15 mL of room temperature HBSS, followed by 20-25 mL of modified Karnovsky’s solution for fluorescence preservation in EM microscopy applications (2.5% Glutaraldehyde, 2.0% PFA, 0.15M Cacodylate buffer, 2mM CaCl_2_, pH 7.2-7.4). The brains were then extracted and post-fixed for 2-24hrs. Following postfixation, the hemispheres were separated sagitally and slices were cut at 500-700μm thickness. The slices were then washed in cold PBS (pH 7.4) 3 x 5 min and immersed in sodium borohydrate dissolved in PBS (10 mg/mL) to quench autofluorescence of glutaraldehyde for 30 min to 1 hr. The samples were then washed once more in cold PBS 3 x 5 min and transferred to glass slides.

### 2-Photon Imaging

The samples were moisturized with one or two drops of ice-cold PBS and covered with Grace Bio-Labs Coverwell imaging chambers. This prevented any shifting in the position of the slice during imaging and further maintained the internal moisture from evaporating during the course of the 2-photon image acquisition. We used a ThorLabs Bergamo II microscope with resonant scanner and a DeepSee Laser with tunable wavelength, using a 25X LWD water immersion objective from Nikon (NA 1.1). We then probed the striatum to find regions where MSNs expressing EGFP could be seen to interact with tRFP-expressing astrocytes. These tended to occur at the boundary of the striatal injection site, as the region surrounding the needle path contained too many labeled astrocyte domains to distinguish which astrocyte was servicing which dendrite.

Multiple ROIs were selected where appropriate and these were scanned at several digital magnification settings, ranging from 1X to capture a general overview of the ROI, up to 8X to record information relevant to individual branches. We used 900-950nm wavelength excitation for green channel and 1000-1050nm for red astrocytes, and all image stacks were recorded at 1280p resolution with z-step size of 0.3 to 1.0μm, depending on the optical zoom and desired detail for that configuration (4X and 8X digital zoom stacks were obtained at 0.3μm z-steps). Following both detailed and overview stack captures, regions were selected for the execution of NIRB protocol (14). The laser was set to 900-950nm and power settings were adjusted to at least 200mW at the back focal plane of the objective. Box scan settings were used with the resonant scanner and small rectangular ablations were made in the vicinity of the ROI for multiple purposes. Larger ablations were made within 300 μm of the ROIs at lower zoom settings (1X or 2X digital zoom), while smaller ablations at higher digital zoom settings were made within 50μm of the desired ROI dendritic branches (**Video 2**). These were closely monitored during generation until the tissue displayed autofluorescence and the markings were clearly visible in transmitted light. The ROI were once more scanned post-branding to record image stacks of the dendritic branches, astrocyte domains, and their relative orientation with respect to the induced ablations. The sagittal sections were removed from the imaging chambers and ROIs were excised with a micro-scalpel under transmitted light. Samples were then immersed in 4% glutaraldehyde and stored for no more than 24hrs before EM treatment.

### EM sample treatment

The samples were prepared in accordance with Mark Ellisman’s lab protocol designed to enhance signal for backscatter electron detection of epoxy embedded mammalian tissue at low accelerating voltages (50). The sample treatment was selected due to its focus in highlighting membrane contrast. All solutions made with double distilled water (ddH_2_O) unless otherwise indicated and all solutions were made fresh for each set of concurrent samples to be prepared. Samples were kept in ice cold fixative (2.5% glutaraldehyde, 2.0% PFA, 2mM CaCl, and 0.15M cacodylate buffer) for 2-3 hours. Samples were rinsed in cold cacodylate buffer containing 2mM CaCl for 3 x 5 min. Right before use, we mixed equal parts of 4% osmium tetroxide and 3% potassium ferrocyanide in 0.3M cacodylate buffer with 4mM CaCl. The tissue was submerged in this solution for 1 hour, ice cold. Samples were rinsed again 3 x 5min with ddH_2_O at room temperature and incubated in thiocarbohydrazide (TCH, 0.22mm Millipore filtered) solution for 20 min at room temperature, followed by 3 x 5min rinse with ddH_2_O. Following rinsing, we incubated with 2% osmium tetroxide in ddH_2_O for 30min at RT. Samples were once more rinsed 3 x 5 min with ddH_2_O at RT and then incubated in 1% uranyl acetate at 4°C overnight. Afterwards, samples rinsed 3 x 5 min with ddH_2_O at RT followed by *en bloc* lead aspartate staining for 30 min at 60C (0.02 lead nitrate in 0.03M lead aprtate, pH 5.5). Samples were whashed once more 3 x 5 min with ddH_2_O and dehydrated in a series of grade ethanol concentrations (20%, 50%, 70%, 90%, 99%, and 100% anhydrous). Each immersion was done 2 x 7 min and then samples were placed in ahydrous ice cold acetone for 10 min, transferred to RT acetone for 10min thereafter. Infiltration with resin was done also with a graded series of Durcupan:acetone (25%, 50%, 75%, and 100%), each for 2-3 hrs at RT with the final step incubating overnight. The next day, samples were immersed in fresh 100% durcupan for 6 hrs and embedded in resin using flat molds to polymerize over 48 hrs at 60C in the oven.

### microCT Imaging (X-Ray)

We used a Versa520 microCT from Zeiss for the x-ray tomography of the embedded tissue blocks. The excess resin was trimmed as much as possible from the boundaries of the blocks and the sample was positioned as close as possible to the x-ray source, to achieve the highest possible resolution. The versa 520 utilizes a cone-shaped source, and the effective voxel size depends on the distance between the detector and the sample. The sample was glued to a holder and scanned twice at 40X magnification. The first scan was done at low voxel resolution to capture the entirety of the sample and pinpoint the location where the large 2-photon ablations could be clearly discerned. The second scan was done at the ROI at 0.6μm voxel resolution, to better resolve the location of the somas within the tissue block and properly overlay the fluorescence dataset using imaging software. The scans were done at 60kV accelerating voltage and 2s dwell time for low resolution scans, or 10s dwell time for high resolution scans of the ROI region.

### Correlation of X-ray and 2-photon datasets

Both X-ray datasets (low and high resolution settings) were loaded onto Avizo Amira (ThermoFisher) and aligned accordingly. Cell somas, 2-photon fiduciary ablations, and blood vessels were used to overlay the 2-photon datasets on the correct region of the x-ray datasets. Once the appropriate location was selected by crossreferencing using all the available x-ray and 2-photon datasets, the block was further trimmed down to just one column of tissue with sides roughly 300-350μm in length (**Figure S1 A**). This was necessary because the microtome blade of the SEM cannot accommodate for consistent surface cuts if the dimensions of the block face exceed 400μm in length. Blood vessels, cell nuclei and induced IR ablations were all used to identify depth, blade angle of attack, and xy coordinates of ROI before selecting trimming region (**Figure S1 B**).

### SBF-SEM imaging

In preparation for SBF-SEM, the trimmed sample was sputter coated with 10nm of gold and inserted into the sample holder of the SBF SEM TeneoVS system from FEI (ThermoFisher). The sample surface was polished at blade speed of 400 mm/s and 100nm thickness until at least 25% of the block surface was revealed. Because the process of SBF-SEM data acquisition is destructive, it is necessary to continuously trim the resin on the sample and estimate the distance to the forthcoming ROI beneath the surface (**Figure S1 B**). The system was then pumped down to vacuum and allowed to stabilize overnight. We then captured a series of preliminary 20-30 slices at 50nm thickness, preferably in regions containing vascular details. (1.78 keV and 10 nA, 2048 by 1768 pixels, 3 μs pixel dwell time, 1000-5000X magnification). ROI specific data was acquired at 40nm cuts. We used 2 energies for true 20 nm axial resolution (1.78 and 2.7 keV, 10 nA current, 2μs dwell time) and corresponding image xy resolution of 6-7nm pixel size, and captured ~200-300 slices per block.

### Data alignment and segmentation

The captured data was aligned and cross-referenced against the microCT scans and 2-photon data prior to segmentation. Once the acquired serial EM block was aligned and guaranteed to contain the ROI of interacting astrocyte and MSN domains, the relevant dendritic branches were manually segmented with the use of Amira software (ThermoFisher Scientific; **Video 1**). Astrocyte structures surrounding the individual dendritic spines were only reconstructed locally, as segmenting the astrocyte for the entire volume was not necessary. For clearly defined regions of analysis containing dendritic spines covered by the overlapping astrocyte domain, the thin and mushroom dendritic spines exhibiting astrocytic engagement were segmented. As a standard, we utilized the same color code for all EM datasets, where the dendritic spine was labeled green, astrocyte in blue, axon terminal in red, and PSD in yellow (**Video 3**). These classes were defined according to the existence of clearly defined PSD structures and proximal astrocyte processes surrounding the synaptic cleft.

### Automated structural analysis from geometry derivation

The automated analytical method used is based on a derivation of the n-dimensional Hausdorff measurements obtained from the interactions of an observed object with respect to a reference object (51, 52), in this case to assess the quantitative information associated with the intersecting planes of the astrocyte process and the PSD (16). We measured the shape and position of the observed astrocyte process (X) with respect to the reference PSD object (Y) at increasing radii *r* from the PSD boundary. Iteratively, we dilate the PSD structure at some distance *r* and investigate the relationship between the PSD-based boundary and the observed object. Here, we measure the section of the astrocyte process that lies within distance *r* from the PSD, denoted Y, where the set of all points within this distance *r* that belong to the astrocyte is called the *r-parallel* set of Y, denoted Y^r^. Therefore, the coordinates corresponding to the astrocyte process (X) object contained within Y^r^, is the intersection of the PSD-derived surface and the astrocyte process at that distance *r*, denoted X ∩ Y^r^.

The interpretation of the intersecting PSD-astrocyte boundary is expanded on **figure 3**, with the corresponding equivalent measurement when the same interaction is observed in 3D parameters. Thus, for any X ∩ Y^r^ where r ≤ 1000 nm from Y, we measured the volume and outer boundary surface area of the intersection boundary parameters of X at distance *r*. Derivation of Hausdorff measurements for intersecting boundaries yielded 2 additional geometric parameters (surface area parallel to reference object, as well as length of intersection perimeter), but these metrics do not translate to intelligible biological analogues, whereas the surface area can be directly understood as the astrocyte membrane and the volume as the internal contents of the astrocyte process. All measurements were normalized for spherical parameters, e.g. volume increase at each r distance was normalized with regards to equivalent total volume of the dilated surface at the same radius, and both normalized and un-normalized results are reported (**Figure 4; Figure S4**). Data was reported for recorded results from an interval of radii representing close proximity to the PSD (25-175 nm from PSD).

### Data Analysis

All statistical comparisons were made with the use of GraphPad Prism 8. Comparisons for non-equal variances, two tail t-test was used for **Figure 2** data. Quantitative results are shown as mean ± S.E.M. and statistical significance was accepted at p<0.05. For **Figure 4**, in order to estimate the statistical significance of the difference between the measurement curves for the WT and R6/2 groups, we implemented the Monte Carlo Permutation test. The distance t’ between the WT and R6/2 mean curves was first measured by the absolute area-under-the-curve. We then collected the graphs from both groups and randomly sampled two new groups from the full collection of curves *n* times, and measured the distance t_1_, […], t_n_. The fraction of t_1_, […], t_n_ < t’ determines the p-value in a stochastic manner (53).

## Resources Table with RRIDs

See Supplemental Information.

## Data and Code Availability

https://qim.dk/software-tools/

https://lab.compute.dtu.dk/QIM/tools/relationalshapemeasure

## Acknowledgments

We thank Emma Mantel for SEM data processing, and the staff of the University of Copenhagen Center for Imaging for technical support. Supported by the Novo Nordisk Foundation and the Lundbeck Foundation.

