## supplemental figures for "Astrocytic engagement of the corticostriatal synaptic cleft is disrupted in a mouse model of Huntington disease"

**This PDF file includes:**

Figures S1 to S4

Table S1

Legends for Movies (Videos) S1 to S4

**Other supplementary materials for this manuscript include the following:**

Movies: Videos S1 to S4

**Figure S1** *Related to Figure 1*

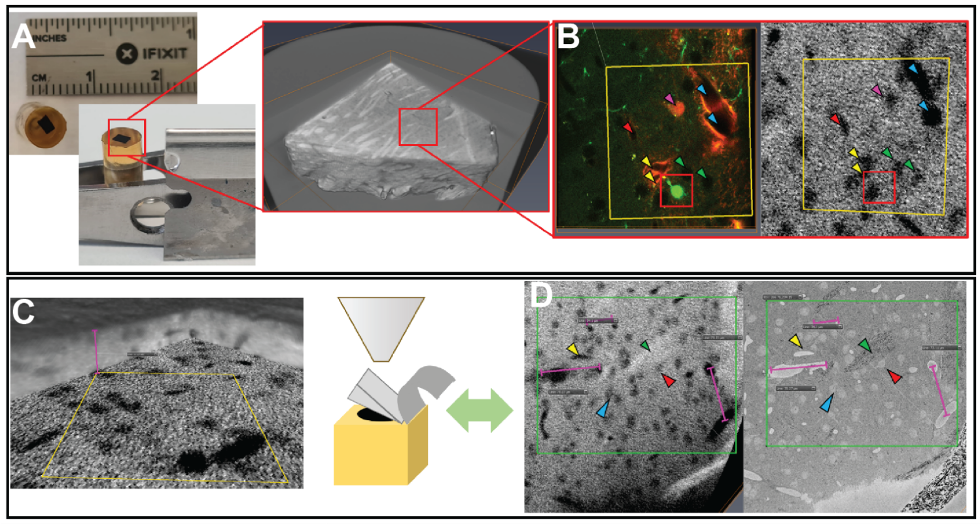

**MicroCT mapping of EM block region containing the desired region of interest**

**(***Related to Figure 1)*

**A-B.** Extracted section of striatum after osmium infiltration and resin polymerization. Sides of block ~500µm, thickness ~600 µm. MicroCT scan of resin block leaves internal structure intact and highlights contrast differences between grey and white matter, as well as vascular cavities and induced ablations, the latter as empty (black greyscale values) **(A).** Comparison of single slice at MSN-astrocyte region of interest (ROI), with highlighted structures relocalized in the microCT scan **(B).** **C.** Use of microCT data to determine location depth of ROI, after cross referencing correct location from 2-photon stack. **D.** Comparison of expected readout from microCT (*left*) and initial surface sections of serial-blockface SEM data (*right*), highlighting induced ablations (*yellow* arrow), myelinated axons (*green* arrow), and corresponding cell soma (*blue* and *red* arrows). Dimensions of measured distances are also shown to match (violet measurements). This confirms expected number of microtome slices remaining to astrocyte-neuron ROI branches, and hence angle estimate at which the diamond blade is cutting the block-face during sample approach. Thus, the microCT data allows us to adjust in real-time the acquisition parameters as the ROI appears in the field of view.

**Figure S2**
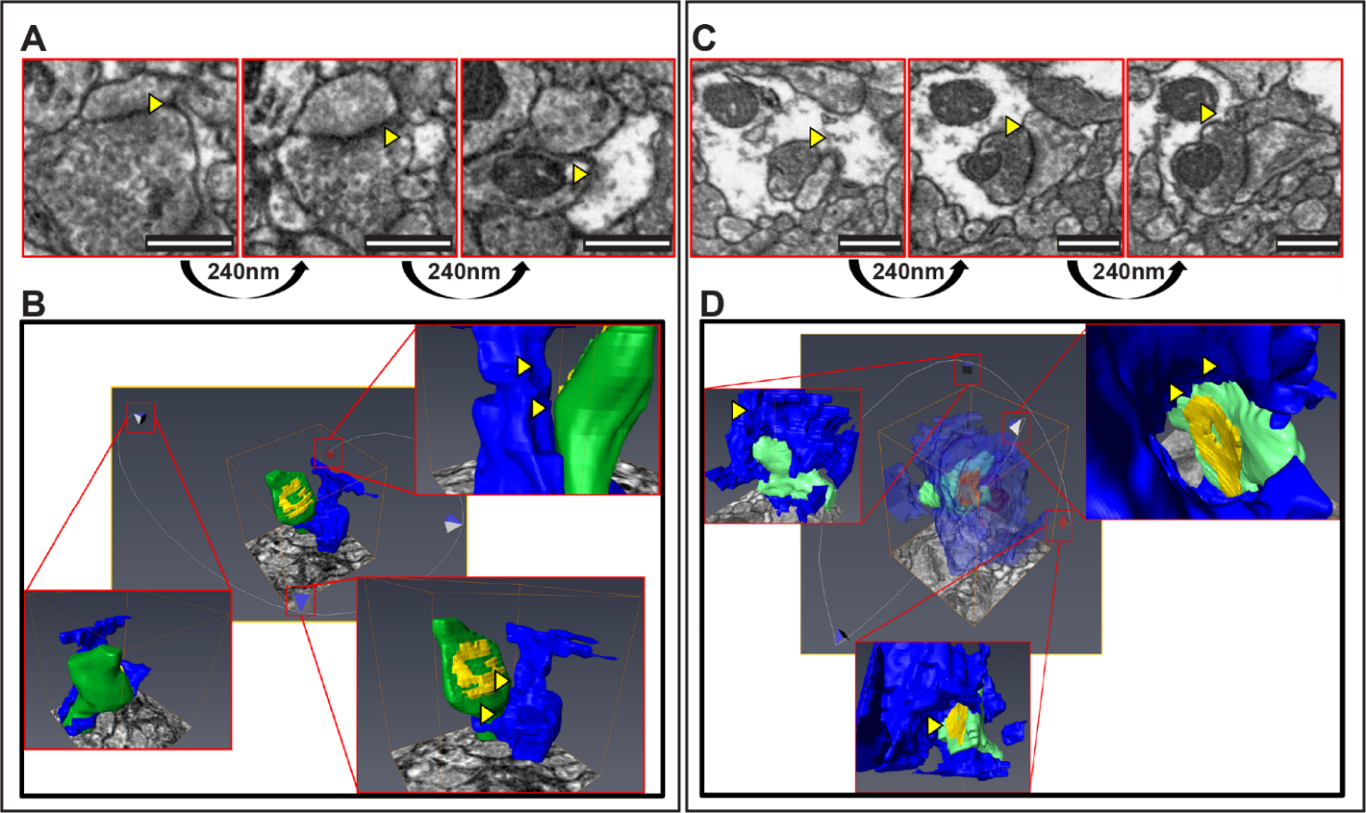

**Morphological variability of synapse-astrocyte interactions (***Related to Figure 4***)**

Three-dimensional reconstruction of spines and astrocytes showing the morphological  variability observed in our datasets. Dendritic spines in *green*, astrocytes in *blue*, and PSDs in *yellow* for the 3D renderings. **A.** Partial astrocyte infiltration into the perisynaptic region of a synapse displaying perforated PSD, as seen in the raw dataset orthogonal to synaptic cleft. **B.** 3D view of thin synapse shown in A, after relevant structures have been segmented using Amira software. Camera path and structures shown at selected vantage points indicating regions of astrocytic engagement, yellow arrows. **C.** Complete astrocytic coverage of a mature synapse with perforated PSD structure. **D.** 3D view of the mature mushroom synapse of panel C, showing a transparent astrocyte structure in the center panel, with the camera path focusing on surrounding corresponding vantage points. Scale: 500 nm.

**Figure S3**

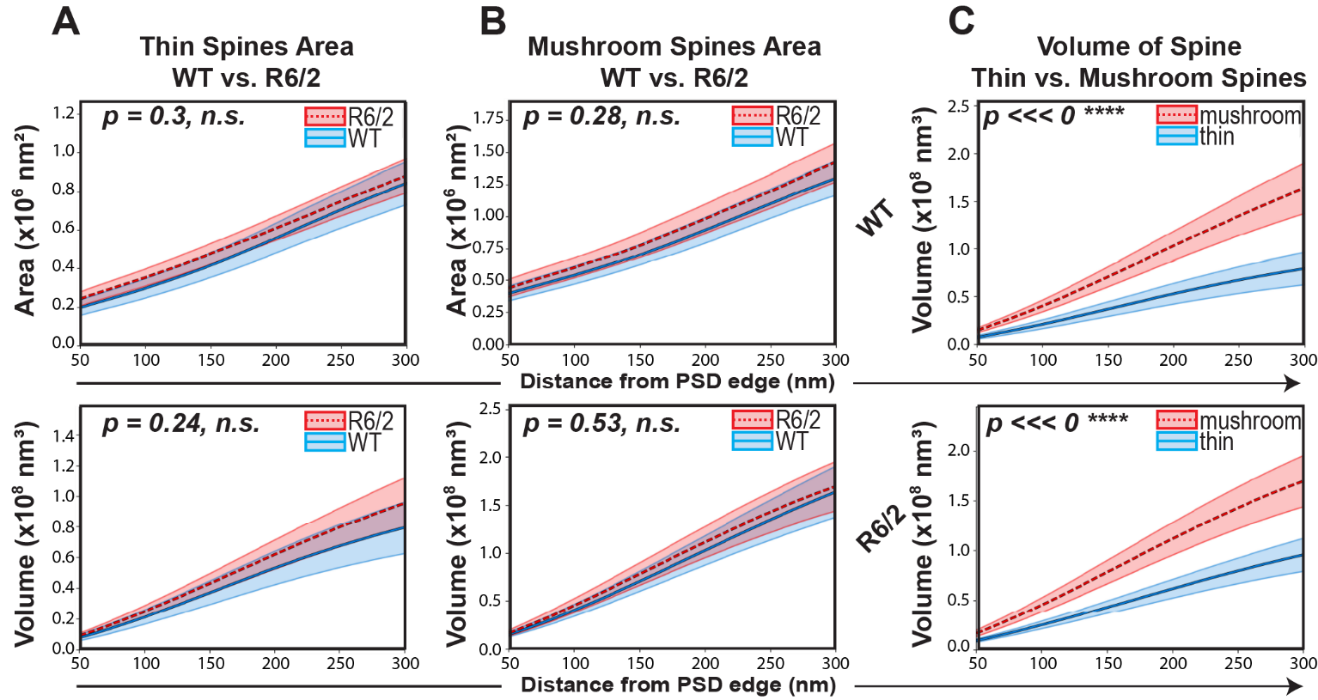

**Comparison of thin and mushroom spines (***Related to Figure 4***)**

Geometric analysis of surface area and volume of cellular components surrounding the PSD, measuring the dendritic spine (post-synaptic terminal) rather than the astrocyte, across both spine classes and disease genotype. **A** (*top panel*) Equivalent area of spine for thin spines across WT and R6/2 MSNs, indicating that the classification used was consistent between groups (p=0.3). Similarly, thin spine volume did not differ between R6/2 and WT mice (*bottom* *panel*; p=0.2).

**B.** Equivalent surface area (top panel) and volume (bottom) of dendritic spines across groups, for synaptic interactions with mature spines (p=0.3) and immature spines (p=0.5), respectively.

**C.** Significantly increased dendritic spine volume was noted, however, for mature mushroom spines in both WT (*top panel*, *** p<0.001) and R6/2 mice (*bottom*, *** p<0.001), indicating that mature spines are larger and more complex than thin/immature spines in both WT and R6/2 MSNs.

*Periphery limits: 50-300 nm from PSD. Thin/Immature spines: WT, 4 mice, n=30 spines; R6/2, 5 mice, n=35 spines. Mature/mushroom spines: WT, 4 mice, n=27 spines; R6/2, 5 mice, n=25 spines. Monte Carlo permutation method, absolute AUC, 95% CI shown.*

Figure S4 *Related to Figure 4*
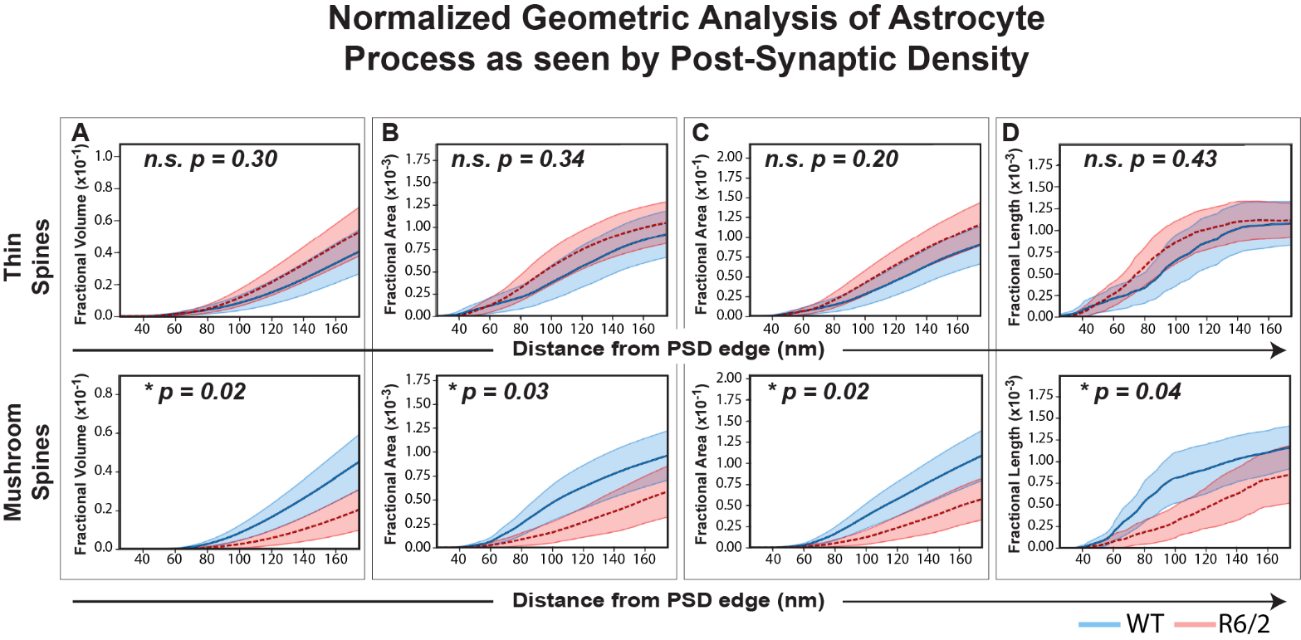

Normalized geometric analysis of astrocytic engagement with corticostriatal synapses

K-cross section, normalized statistics for astrocytic presence within the perisynaptic region encompassing 25 nm to 175 nm from the PSD edge. Data represent a description of the geometry of the astrocytic processes invading the perisynaptic volume, as seen from the PSD.

A. Thin/immature spines in R6/2 mice (*top* panel) showed no change in the fractional proportion of perisynaptic volume occupied by astrocytic processes, relative to wild-type control synapses (p=0.3). In contrast (*bottom* panel), the perisynaptic space surrounding (dilated from) mature/mushroom spines exhibited significantly decreased astrocytic volume, relative to wild-type (*p=0.02), indicating significant astrocytic atrophy in the periphery of the PSD for R6/2 mice.

B. Similarly, astrocytes associated with thin/immature R6/2 spines (*top* panel) showed no change in their fractional surface area (p=0.34), while (*bottom*) astrocytes associated with mature/mushroom R6/2 spines display decreased fractional surface area (*p=0.03).

C. Astrocytes associated with thin/immature spines (top panel) showed no change in their fractional perpendicular surface area as a function of disease (p=0.2). In contrast, R6/2 astrocytes surrounding mature/mushroom spines (*bottom*) displayed significantly decreased fractional perpendicular surface area (*p=0.02).

D. Astrocytes associated with thin/immature spines in R6/2 mice (top panel) were no different from wild-type controls in their perisynaptic fractional contour length (p=0.43); in contrast, astrocytes engaged with mature mushroom spines of R6/2 MSNs showed significantly decreased contour lengths (*bottom*) (*p=0.04).

*Thin/Immature spines: WT, 4 mice, n=30 spines; R6/2, 5 mice, n=35 spines. Mature/mushroom spines: WT, 4 mice, n=27 spines; R6/2, 5 mice, n=25 spines. Monte Carlo permutation method, absolute AUC, 95% CI shown.*

**Table S1**

**RESOURCES TABLE**

| Reagent or Resource | Source | Identifier |
| --- | --- | --- |
| Chemicals |  |  |
| Sodium borohydrate | Merck, Sigma-Aldrich | Cat# 452882-100G |
| Hanks Balanced Salt solution +/+ | ThermoFisher Scientific | Cat# 24020091 |
| Modified Karnovsky (2.5% Glutaraldehyde, 2.0% Paraformaldehyde) | Electron Microscopy Sciences (EMS) | Cat# 15732-10 |
| 10X PBS | EM Sciences (EMS) | Cat# 19342-10 |
| Xylocaine (lidocain) | Accord Healthcare Limited | Cat# 146901 |
| Carbprofen (Rimadyl) | Zoetis Finland OY | Cat# 102799 |
| 4% osmium tetroxide | EM Sciences (EMS) | Cat# 19170 |
| Potassium ferricyanide | EM Sciences (EMS) | Cat# 20150 |
| Uranyl acetate | EM Sciences (EMS) | Cat# 22400 |
| Lead nitrate | EM Sciences (EMS) | Cat# 17900 |
| Aspartic acid | Merck, Sigma-Aldrich | Cat# A9256-100G |
| Thiocarbohydrazide | EM Sciences (EMS) | Cat# 21900 |
| Ethanol (95%) | Merck, Sigma-Aldrich | Cat# 51976-500ML-F |
| Ethanol (100% anhydrous) | EM Sciences (EMS) | Cat# 15058 |
| Embed-812 | EM Sciences (EMS) | Cat# 14121 |
| Propylene oxide | EM Sciences (EMS) | Cat# 20401 |
| Recombinant DNA |  |  |
| RV-dG-EGFP | Viral Vector Core - Salk | N/A |
| Lenti-GFAP-tRFP-Mir124T | This paper | N/A |
| Software |  |  |
| Amira | ThermoFisher Scientific | RRID:SCR_007353 |
| NIS-Elements | Nikon Instruments Inc. | RRID:SCR_014329 |
| ImageJ | NIH ImageJ | RRID:SCR_002285 |
| Python | Python Software Foundation | RRID:SCR_008394 |
| C++ | isocpp.org | N/A |
| Prism | GraphPad Prism | RRID:SCR_002798 |
| Animals |  |  |
| Mouse: B6.129S7-*Rag1^tm1Mom^*/J | Jackson Laboratory | RRID:IMSR_JAX:002216 |
| Mouse:B6CBA-Tg(HDexon1)62Gpb/3J (CAG 120±5) | Jackson Laboratory | RRID:IMSR_JAX:006494 |
| Other |  |  |
| Coverwell imaging chamber, 0.6mm high | ThermoFisher Scientific | GBL635051-40EA |
| Leica VT1000 S Vibratome | Leica Microsystems | RRID:SCR_016495 |
| Zeiss X-Radia XRM520 Versa | Carl Zeiss AG | N/A |
| Leica Ultramicrotome ARTOS 3D | Leica Microsystems | RRID:SCR_020226 |
| ThorLabs Bergamo II | ThorLabs | N/A |
| MaiTai DeepSee DS+, pulsed laser | Spectra Physics | N/A |
| Nikon CFI75 Apo 25XC W | Nikon Instruments Inc. | MRD77220 |
| VolumeScope 2 SEM | ThermoFisher Scientific | RRID:SCR_019913 |
| Nikon SMZ1270i stereomicroscope | Nikon Instruments Inc. | RRID:SCR_020328 |

**Movies**

**Video 1**

**Correlative microscopy, 2 photon and X-ray imaging for ROI definition**

*Related to Figure 1*

X-ray datasets of samples were used as a main navigation tool, whereby the higher resolution 2 photon datasets from preceding captures are overlayed onto both fiduciary marks and the microvasculature, to determine the correct x-y-z coordinates of the prospectively defined neuron-astrocyte region-of-interest (ROI), for subsequent serial SEM imaging. Note that the reconstructed MSN details the difference between the 2 photon-segmented and serial EM datasets.

**Video 2**

**Relocalization of EM stack, after X-ray and 2 photon correlated light EM (CLEM)**

*Related to Figure S1*

This video presents the EM environment with iteratively higher magnification scans. Multiple EM stack correlations are necessary, since serial block-face SEM is a destructive imaging process, and there is only one opportunity to capture each desired ROI stack. The reconstructed medium spiny neuron branch shown here is the same as that in video 1, with dendritic spines added to show the neuron-glia environments selected for analysis.

**Video 3**

**Reconstructed neuropil volume of striatum and perisynaptic astrocyte**

*Related to Figure 2*

This selected subvolume of striatum shows the density of the segmented environment. The camera path is focused on a single synapse, and shows a prototypic example of astrocytic coverage of the perisynaptic region. Multiple views reveal the complex association of astrocytes with both the pre- and post-synaptic components of mature, mushroom-like synapses, the proximal elements of which were used for the analysis strategy described in Figure 2 (see segment with opaque and transparent overlay of astrocyte, at 00:22:00).

**Video 4**

**Identification and characterization of perisynaptic morphologies**

*Related to Figures S2 and S3*

We observed a variety of configurations of astrocytic engagement with MSN spines, with a spectrum of astrocytic infiltration patterns observed at each analyzed level of dendritic spine maturity. These examples focus on the prototypic engagement of mature spines in WT mice, in which tight astrocytic sequestration of the perisynaptic space was typically observed.
